# Nanoparticles of bioactive natural collagen for wound healing: Experimental approach

**DOI:** 10.1101/2023.02.21.529363

**Authors:** Manal Shalaby, Ahmed .Z Ghareeb, Shaimaa M. Khedr, Haitham M. Mostafa, Hesham Saeed, Dalia Hamouda

## Abstract

**Introduction:** Both developing and developed nations have made the creation of innovative wound-healing nanomaterials based on natural extracts a top research goal. The objective of this research was to create a gel containing collagen nanoparticles and evaluate its therapeutic potential for skin lesions.

**Methods:** Collagen nanoparticles from fish scales were produced for the first time using desolvation techniques. Using Fourier transform infrared spectroscopy (FTIR), the structure of the isolated collagen and its similarities to collagen type 1 were identified. The surface morphology of the isolated collagen and its reformulation into nanoparticles were examined using transmission and scanning electron microscopy. Human skin fibroblast cells were employed to examine the cytotoxicity of the nanomaterials, and an experimental model was used to evaluate the wound healing capability.

**Results:** Collagen nanoparticles formulation was confirmed using FTIR, SEM and TEM analysis. Cytotoxicity studies demomstrated that the manufactured nanoparticles have minor toxicity at high concentrations on human skin fibroblast. Histological investigation proved that the fabricated fish scale collagen nanoparticles promoted the healing process in comparison to the saline group.

**Conclusion:** The fabricated product is a highly influential wound healing product that can be applicable for commercial use. The nanoscale size of collagen nanoparticles, make them interesting candidates for biological applications.

**Key Summary Points:**

- The goal of this research was to create natural, effective wound remedies that could lower health-care costs while also providing pain relief and, ultimately, effective scar repair.
- Collagen nanoparticles can be synthesized from fish scale utilizing various nanotechnology-based approaches to stimulate skin cell proliferation and promote wound healing.
- Collagen nanoparticles have a rough surface, have a negative potential, and can be used for drug delivery and wound healing.
- Histological and macroscopical analysis showed that the synthesized nanoparticles promoted faster wound healing.

## Introduction

Millions of individuals throughout the world suffer from chronic wounds, which provide a therapeutic challenge that could be solved by the development of better dressings that stimulate tissue regeneration^1^. High viscoelastic, biodegradable, and hydrophilic biomaterials used in wound dressing induce angiogenesis, decrease inflammation, regulate cell adhesion, and promote the growth of keratinocytes and fibroblasts^2, 3^. Interestingly, collagen has been proposed as an ideal candidate for wound dressings. Collagen promotes the development of the numerous cell layers found in the skin, which enhances wound healing. Collagen-based dressings have the potential to reconstruct the intricate collagen architecture of original tissue ECM collagen and ECM-associated secondary components such as laminin, fibronectin, and glycosaminoglycans^5^.

Many biomaterials designed to stimulate skin cell proliferation could be produced as nanoparticles utilizing various physicochemical techniques, allowing for the modification of this biomaterial ^6^. Due to their nanosize and high surface area-to-volume ratios, innovative biomaterial nanoparticles of natural origin have been used in wound treatment ^7^. In an effort to create permeable oxygen-exchange wound healing dressings that do not provoke antigenicity, research on nanomaterials has produced a number of products. In this regard, collagen has been marketed as an attractive natural and safe substitute for drug delivery and therapeutic applications^8^.

Collagen could also act as a promising drug delivery carrier as a result of the existence of the ionizable group such as amino, phenol, guanidine, and imidazole ^9^. Unfortunately, due to its large Mw of about 300 kDa, native collagen is unable to permeate the skin’s SC^10^. While collagen applications in wound healing have been broadly utilized, collagen nanomaterials reports are insufficient, as the pure type-1 collagen from animal sources is expensive and has different forms.

For medical purposes, various forms of collagen were separated from marine sources, and the recovered protein was assessed using physicochemical characterization. Wound contraction and histological examinations were also utilized to test the efficacy of extracted collagen for wound healing. Assessments showed that collagen dressings enhanced wound resolution and closure. According to the findings, isolated fish scale collagen might be a promising alternative for wound healing^11^.

Due to their high loading capacity, successful intracellular drug delivery, efficient cellular absorption, and sustained release in cytosols with efficient biodegradation in endosomes/lysozymes, nano-carriers were proposed as potential therapeutics. Surface roughening of nanoparticles has also been established as another method for increasing biomolecule loading capacity and cellular absorption efficiency ^12, 13^.

Herein, we report a study on the preparation method and transformation of collagen into a new nanomaterial that influences cell propagation, movement, stratification, and ECM deposition, to enable the development of substitute skin tissues and wound dressings. The ARRIVE guidelines were used to conduct this investigation.

## Design and Methodology

All experiments were carried out according to the relevant guidelines and ethical regulations of the Local Ethical Committee, City of Scientific Research & Applied Technology.

### Isolation of collagen from fish

Fish scale collagen (FSC) was extracted from Tilapia scales obtained from sea capture thoroughly washed in flowing water as previously described^11,14^. The scales were then properly cleaned with distilled water, dried and kept at **-**25°C until they were utilized.

The dried scales (5.0 g) were treated with 0.1 N NaOH for 3 days, changing the solution every day. Next, fish scales were cured with 0.5 M acetic acid for 3 days, and the extract was recovered by centrifugation at 50 000 g for an hour.

The salt of NaCl was added gradually to a final concentration of 0.9 M, the supernatants were pooled. To eliminate any salt, the pellet was rinsed and reprecipitated three times with pure water. The pellets were lyophilized after being suspended in 0.5 M acetic acid. Using a personal mill, the lyophilized samples were ground to a powder and sieved with a mesh (0.15 mm) sieve.

### Spectrum Analysis

Shimadzu FTIR 8400S, Japan, was used to perform FTIR spectroscopy on a freeze-dried collagen sample. A sample of 10 mg was combined with 100 mg of dry potassium bromide (KBr) and compacted to form a disc with a diameter of 10 mm. The spectrum peak was scanned between 4500 and 500 cm^-1^.

### SEM Analysis

The microstructure of fish collagen and fish scale collagen nanoparticles were examined using SEM. The lyophilized collagen sample was punched and attached to an adhesive carbon stub. A tabletop SEM (JEOL 6340, Japan) was used for imaging at a voltage of 15 kV.

### Isolation and Purification of Collagen nanoparticles

The desolvation method was used for the preparation of collagen nanoparticles that was first employed by Marty et al.^15^. This method employs the addition of absolute ethanol drop by drop to reach 50 ml as a desolvation factor to 50 ml of the collagen solution (0.5 g in 50 ml of 0.1 M acetic acid). This would alter the tertiary structure of collagen with the subsequent formation of collagen nanoparticles. Lastly, 500 µl of glutaraldehyde was added as a cross-linking material.

### Characterization of Nile fish scale-based collagen nanoparticles

Light microscope, SEM, TEM, and DLS analyses were performed to characterize the formed Nile Tilapia fish scale-based collagen nanoparticles. For a better dispersion, the solution of Nile Tilapia fish scale-based collagen nanoparticles was sonicated for 5 min to prepare the TEM sample.

The DLS technique was used to determine the size distribution and ζ potential of the produced Nile Tilapia fish scale-based collagen nanoparticles. Measurements were carried out on a Malvern Zetasizer (Malvern Instruments Corp., Malvern, United Kingdom) in solutions of pH=5. All samples were diluted with Millipore-filtered (MF-Millipore™ Membrane Filters) deionized water to an appropriate scattering intensity.

### Collagen nanoparticles cellular treatment for Human skin fibroblast cells (HSF)

Human fibroblast cells were obtained from the BJ-5ta (ATCC® CRL-4001TM) cell line and cultured in D-MEM media supplemented with 10% FBS in 25 cm^2^ flasks at 37 °C and 5% CO_2_. Freeze-dried collagen nanoparticles were dissolved in D-MEM cell culture supplemented medium applied to cells at concentrations of 0.1, 1, 10, and 200 g/mL. Cells were washed twice with a sterile solution of phosphate buffered saline (PBS) and collected for analysis at varied recovery periods (0, 24, 48, and 72 h). Complete media was used as a blank, whereas the negative controls in these studies were untreated cells.

### Preparation of collagen nanoparticles gel

Collagen nanoparticles 0.1 g was collected by centrifugation at 6000 rpm for 30 minutes with 1 ml of a platelet-rich plasma separation gel. The platelet-rich plasma separation gel is created from a resin that is made up of the following fundamental components in weight order: With a specific weight range of 1.068 to 1.080 at 25°C, 120 parts butyl acetate, 75-80 parts phenethylene, 20-25 parts butyl acrylate, 5-12 parts acrylic acid, 2-2.5 parts azodiisobutyronitrile (AIBN), and 0.5-0.8 parts dodecyl mercaptan Thereafter, the collagen nanoparticles gel was collected for further testing^16^.

### *In vivo* experimental study

The current research study was authorized by the board of the Institutional Animal Care and Use Ethics Committee of City of Scientific Research & Technology Applications (Application approval No. 53-3V-0122). The experimental research protocol was employed at the Pharmaceutical & Fermentation Industries Development Center and endorsed by the central committee. This study also followed the procedures established in the guide of the American Guide for the Laboratory animals Care & Use for the maintenance, management, sedation, analgesia, anesthesia, and euthanasia of laboratory animals.

Fifteen healthy indigenous rabbits (5-6 months old and weighing 1.5-2 kg) were used for the research. Animals’ groups were housed in isolated stainless-steel cages with regulated light and ambient temperature on a 12-hour light/dark cycle. Animals were clinically healthy and housed at the Vivarium of the laboratory animal unit, Preclinical Studies Department, Pharmaceutical & Fermentation Industries Development Center (PFIDC), Borg El Arab, Alexandria.

All animals were numbered and weighed before being separated into three groups of five animals each: Group (I) acted as the vehicle control and received saline therapy, Group (II) received PRP gel treatment, and Group (III) received gel containing fish collagen nanoparticles treatment.

I/M injections of 2% Xylazine (1 mg/kg b.wt) and Ketamine HCl (0.5 mg/kg b.wt) were used to sedate the rabbits. Each animal was separated, fasted, and given a broad-spectrum antibiotic, Azithromycin 50 mg/kg, once before surgery and once per day in drinking water for three days afterward. To reduce pain stress on the animal, oral acetaminophen 100 ml drinking water was given for 5 days after surgery.

The surgical region was shaved, and 4% alcohol-based iodine was applied as an antiseptic to the surgical site. Lidocaine 5% local analgesic sub-cutaneous injection at the incision site was applied. At the center of the shaved area on their upper backs, sterile surgical scissors were used to create a scar with a circular diameter of 2.5 cm and a full thickness open wound excision to the extent of subcutaneous tissue. For 21 days, all animal groups were maintained in their cages separately.

### Wound Closure Rates: Planimetry Analysis

On postoperative days 0 to 21, a digital camera was used to take pictures of the experimental animals’ scars in comparison to a metric ruler. The assessment of calibrated wound closure in a two-dimensional plane and its measurement was defined by the appearance of wound edge closure and wound surface re-epithelialization. Four individual photo-micrographic measurements were taken of each rabbit.

### Standard histology of the wounded skin

Sodium pentobarbital was injected intravenously into the ear vein for the animals’ euthanasia before sampling. Sharp dissection was used to obtain a full thickness skin flap tissue specimen from the operation site. Sections were then fixed using 10% formalin with subsequent paraffin processing. Serial sectioning was performed for paraffin sections harvested on Days 10 and 21, and H &E staining was performed for representative sections.

## Results

### FTIR

The secondary structure of collagen found in fish scales was examined using FTIR (Fig. 1). In FTIR spectra between 450 and 4000 cm^-1^, the primary absorption bands of amides A (3427 cm^-1^), B (2937 cm^-1^), I (1651 cm^-1^), II (1545 cm^-1^), and III (1244 cm^-1^) could be detected ^17^.

**Figure (1):**
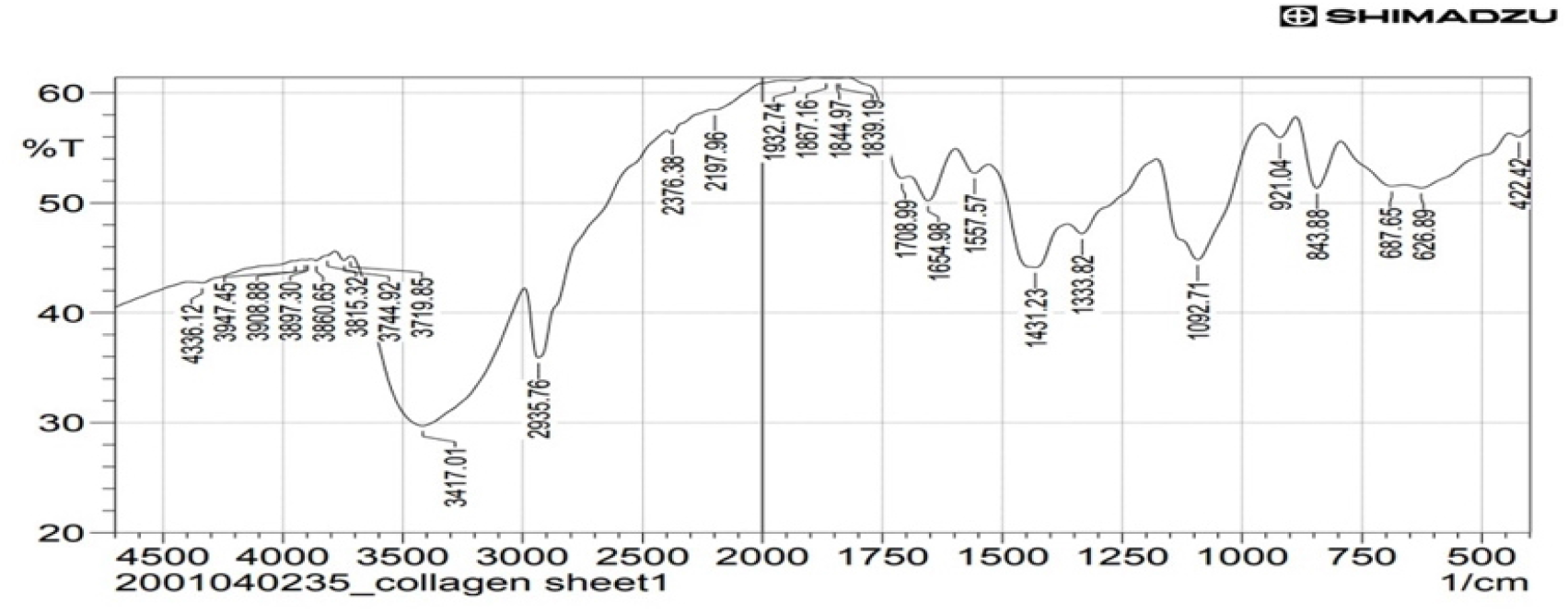
FT-IR spectrum of collagen isolated from Tilapia fish

### Collagen fibers modulation for nanoparticles formation

The purified collagen was examined with optical microscopy to determine the presence of fibrils collagen’s structure (Fig. 2a), which was curbed to tiny spheres upon desolvation (Fig. 2b). The creation of nanoparticles could be indicated by the collagen solution having a milky appearance after the slow addition of ethanol. The effect of temperature and pH on nanoparticle size has been investigated; showing that the optimum temperature was 37°C while the optimum pH was 5.

**Figure (2):**
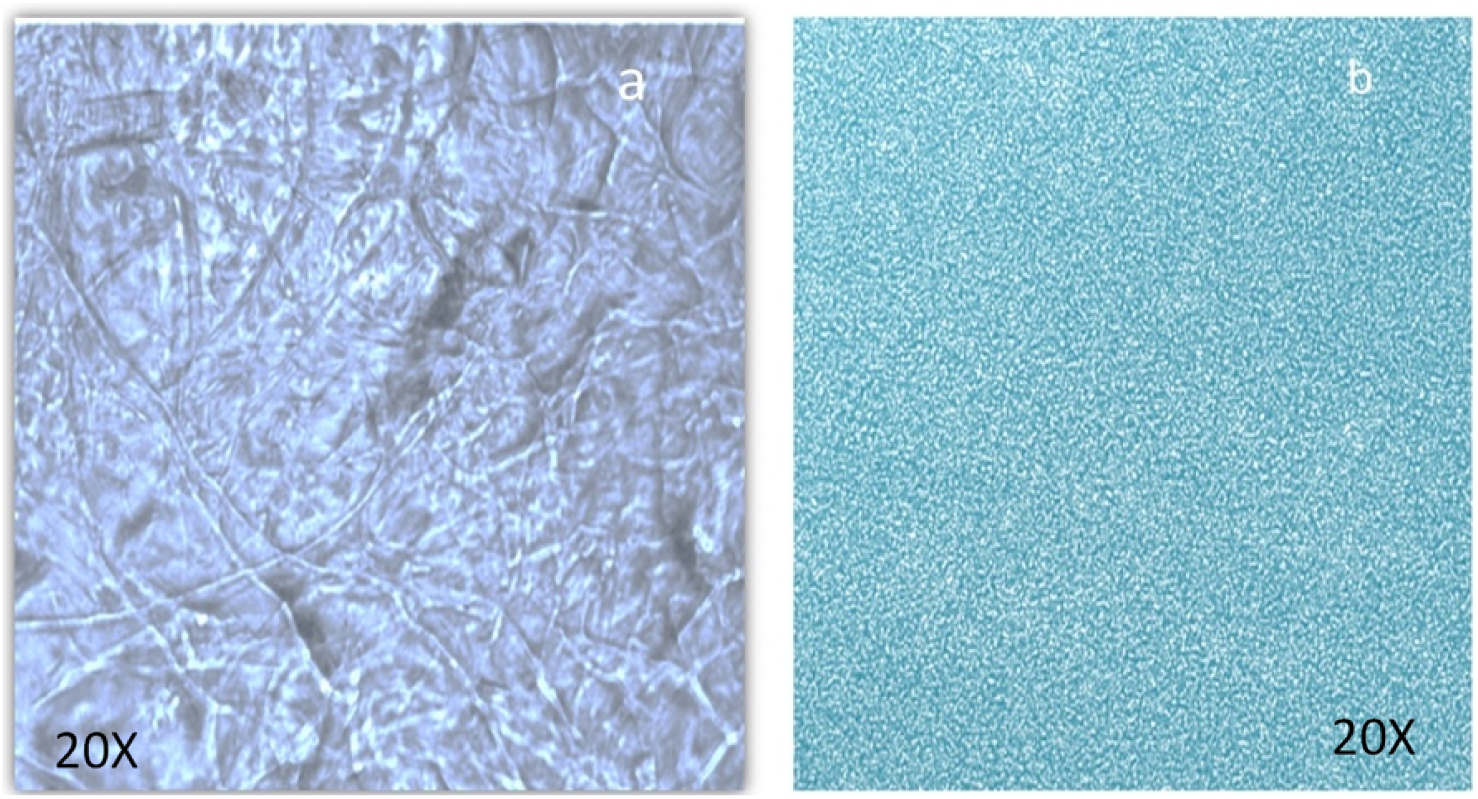
Light microscopy images of collagens fibers prepared from the scales of Tilapia (a) collagen nanoparticles prepared using desolvation-based method (b).

### Scanning electron microscopy (SEM) analysis of nanoparticles

SEM microscopy 370X revealed the fibrillar structure of collagen (Fig. 3). The size of the articulated nanoparticle is between 100 and 350 nm at a magnification of 20000X (Fig. 4)..

**Figure (3):**
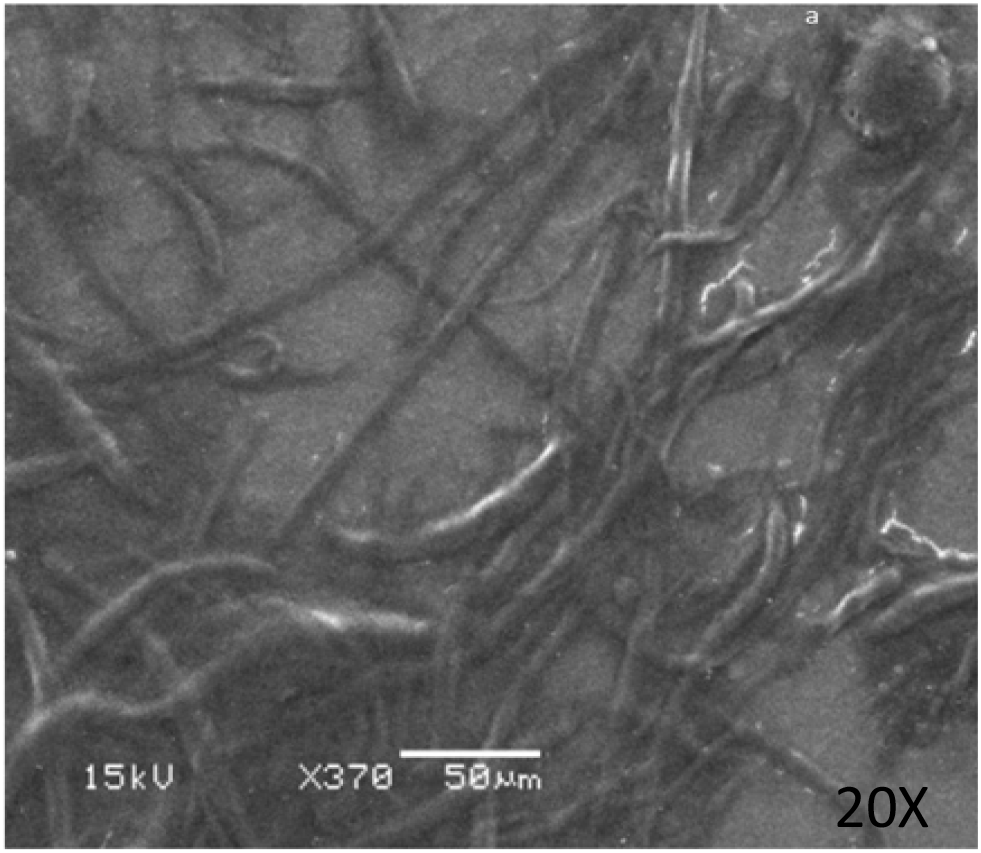
SEM observation of fish scale collagen (X370) showing the fibrillar structure of collagen.

**Figure (4):**
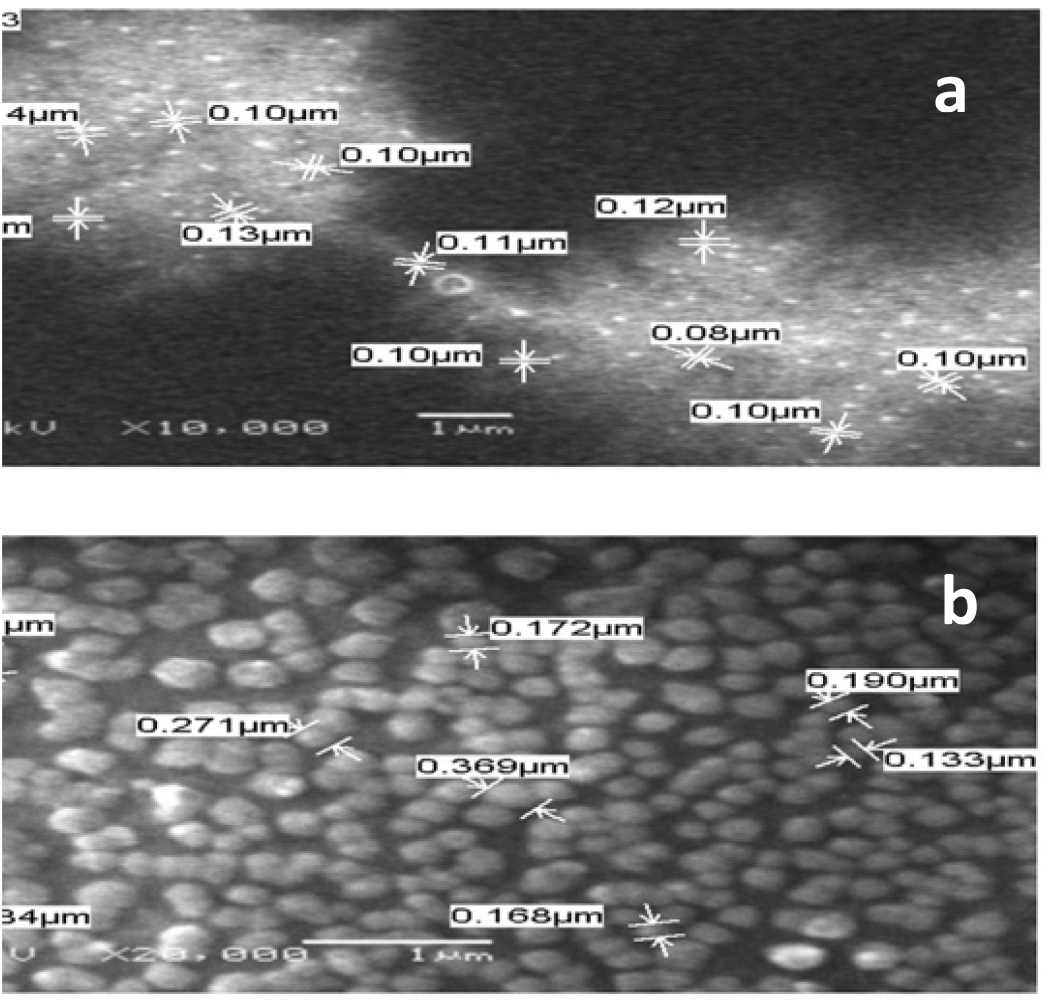
SEM observation of collagen nanoparticles showing an analysis of particle size at different magnifications;10,000 X (a) and 20,000 X (b).

### Transmission electron microscopy (TEM)

TEM images of collagen nanoparticles are presented in Fig. 5, showing that nanoparticles are spherical in structure, having a size of 7.67, 8.57 and 11.71 nm. The surface of most of the nanoparticles is decorated with what seems like a protein shell, forming a biologically active protein corona.

**Figure (5):**
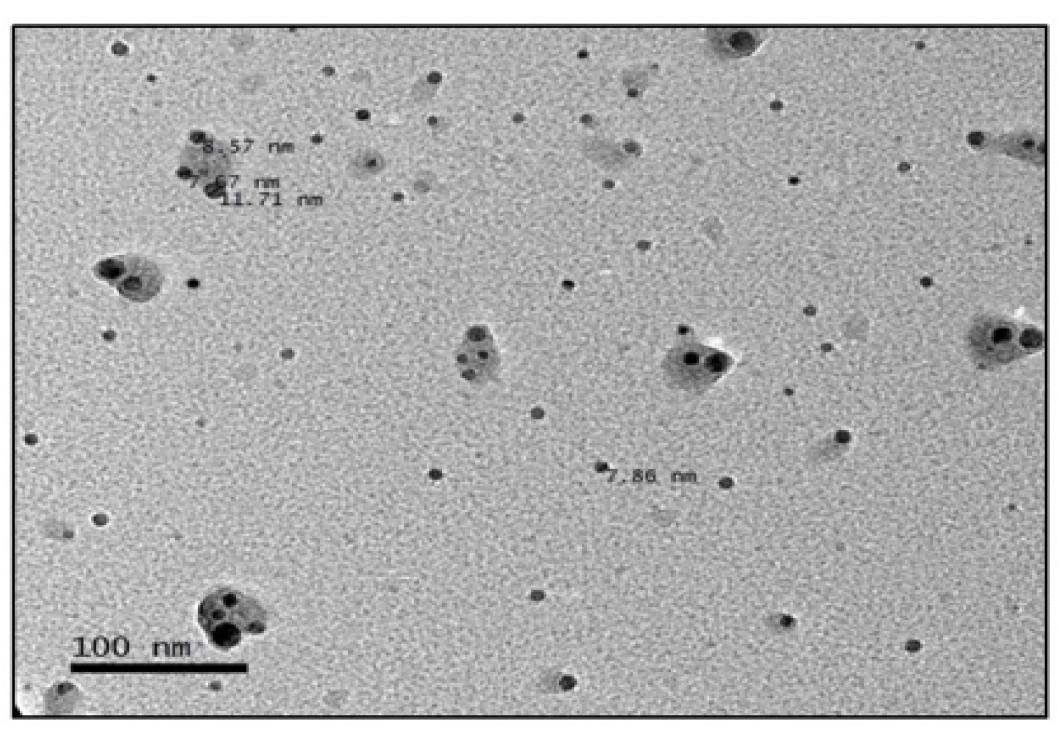
TEM observation of collagen nanoparticles showing an analysis of particles shape and distribution.

### Determination of collagen nanoparticles dimensions

Collagen nanoparticles obtained were polydispersed in nature, with an average <u>diameter</u> of 182 nm and a comparable average <u>zeta potential</u> value of -17.7 mV (Fig. 6).

**Figure (6):**
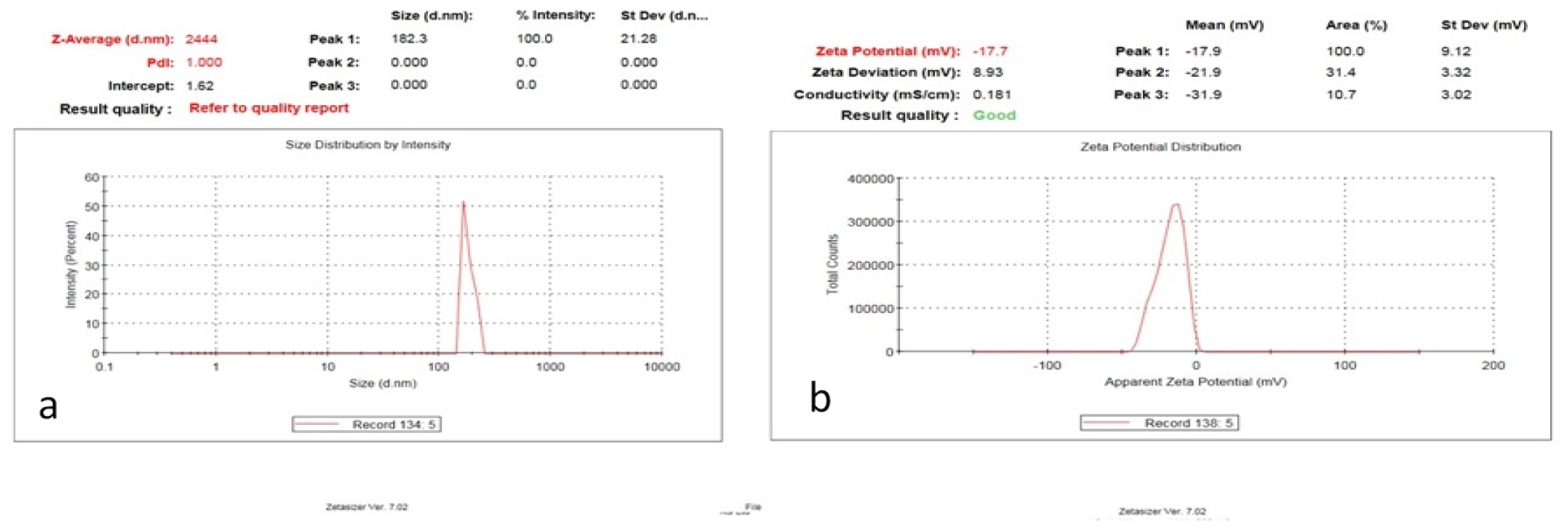
DLS analysis of collagen nanoparticles showing particle sizes (a) and particle potential (b).

### Human skin fibroblast containing collagen nanoparticles

HSF in its natural state is composed of polygonal, adhering cells that form a confluent monolayer (Fig. 7c). Our results showed that treating fibroblast cells with collagen nanoparticle concentrations less than 64 µg/ml for 24 hours resulted in no or very mild morphological alterations, indicating that the cells were alive (Fig. 7). Treatment of fibroblasts with larger quantities of collagen nanoparticles (64 µg/ml), on the other hand, dramatically increases the refractivity of the cell nucleus. Treatments with concentrations of 64 µg/ml and higher demonstrated morphological changes accompanied by increased cell-to-cell contact, implying that the cells started to separate from the base. There were also more cells with visible multinuclear, abnormal vacuoles, or particles in cellular plasma, as well as an increase in elongated cell numbers and visible filopodia.

**Figure (7):**
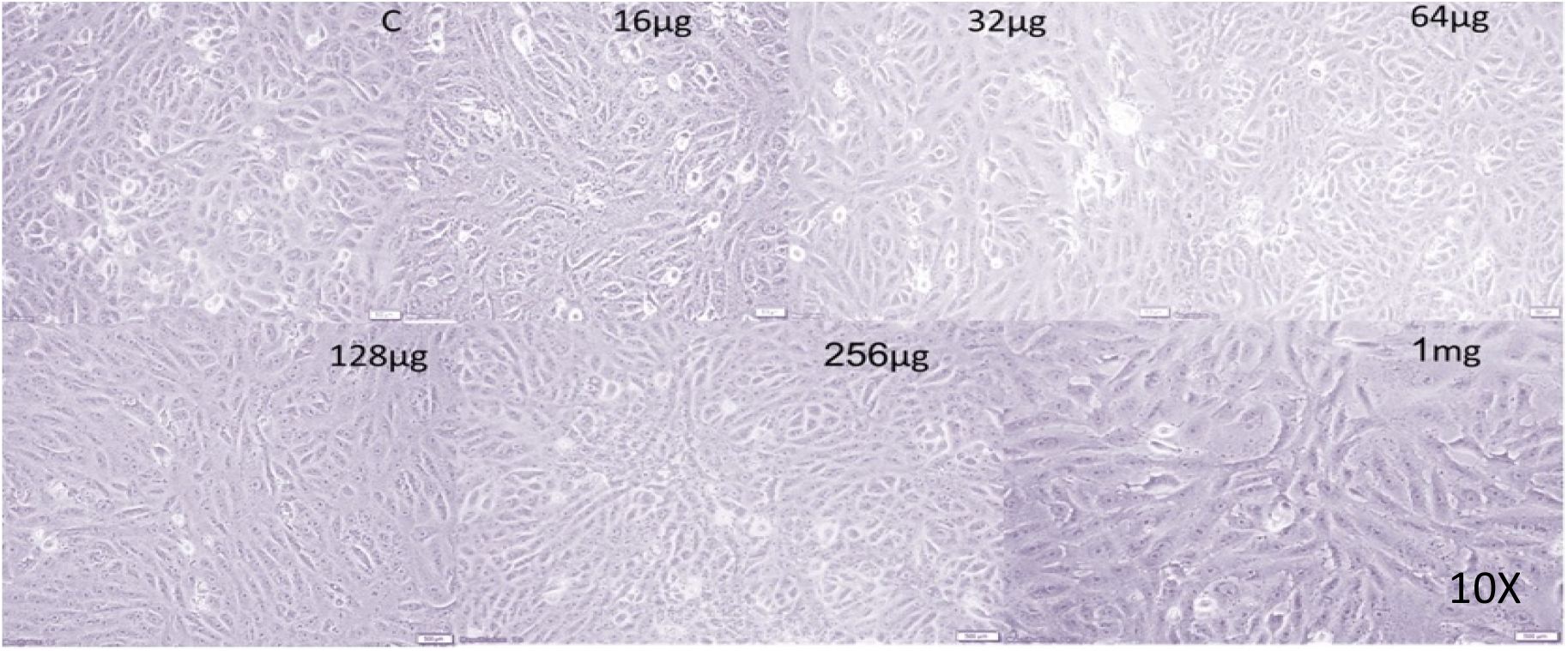
Light microscope image (10x) of human skin fibroblast cells (HSF) subjected to different concentrations (16 µg, 32 µg, 64 µg, 128 µg, 254 µg and 1 mg) of collagen nanoparticles.

### Testing the wound healing potential of collagen nanoparticles

#### *In vivo* wound healing assessment

In comparison to saline, PRP gel (G2) and gel mixed with collagen nanoparticles (G3) demonstrated effective wound sealing within 7–14 days of damage, for quicker and better re-epithelization. It also improved wound closure and had no contamination complications during the healing process (Fig. 8).

**Figure (8):**
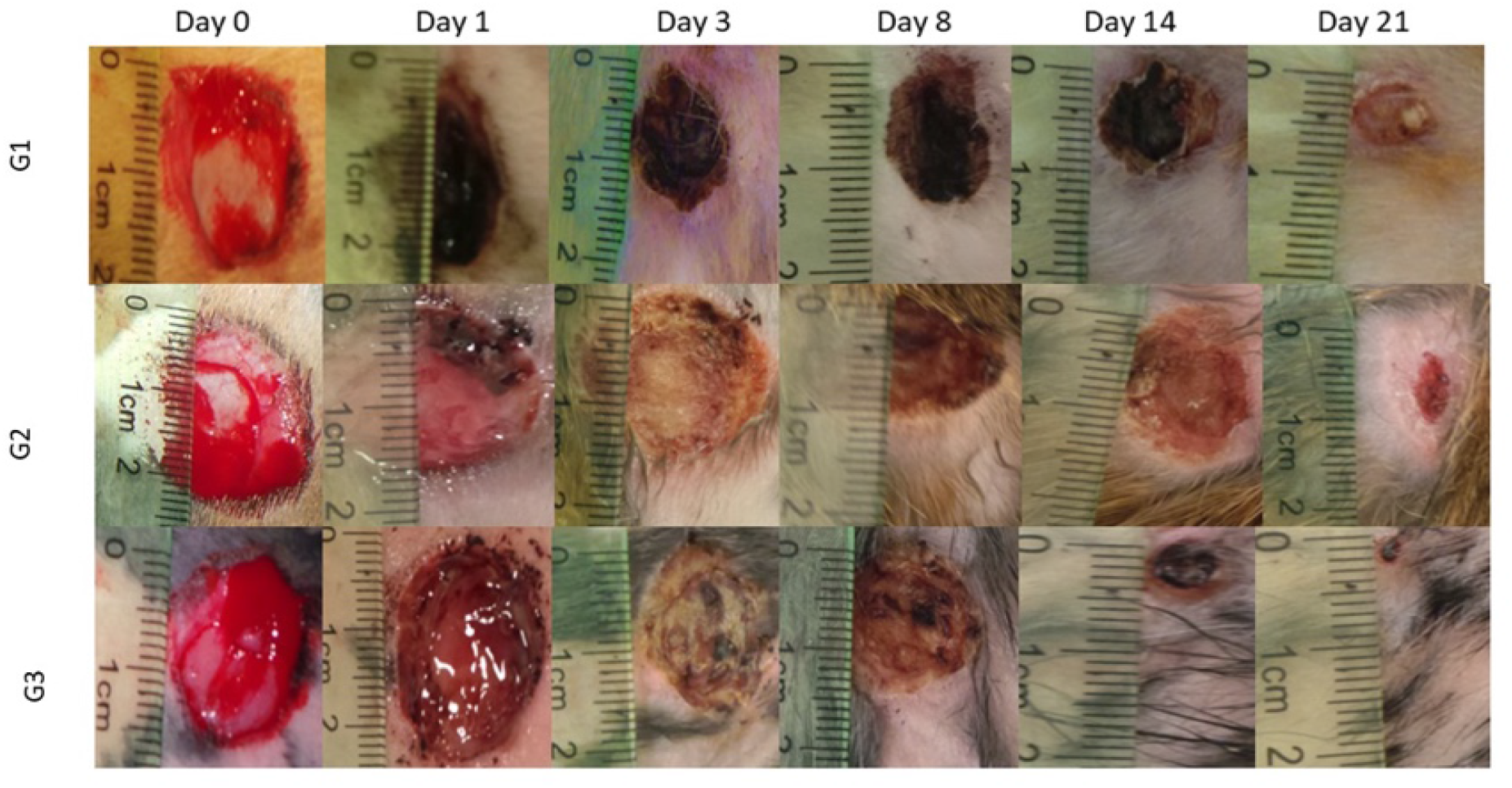
Digital photos showing macroscopic wound size and condition compared with an initial value at day 0 where (G1) saline control group & (G2) represents the group treated with PRP gel and (G3) is the group treated with gel mixed with collagen nanoparticles.

The macroscopic examination of the wounded area (Fig. 8) aimed to detect probable erythema near the lesion, the presence of extreme exudate, and the preservation of wound humidity. In comparison to saline wound healing, collagen nanoparticles restored the injured tissues. Collagen nanoparticles-based dressing were demonstrated to be suitable for dry wound cleaning and facilitated autolytic debridement. This dressing was permeable, did not react with the injured tissue, left a little residue on the wound and aided in wound re-epithelialization. On the third day, all wounds had shrunk in comparison to the operation day; nevertheless, variations could be seen as compared to the saline group gel (G2); in collagen nanoparticles groups (G3), the wound begins to heal up with new tissue, and new skin forms over this tissue. The margins of the incision in collagen nanoparticles groups (G3) draw inward as it heals, and the wound shrinks, displaying a crust all around the wound with granulated tissue detected, suggesting that it had dried.

All important elements of the wound assessment process; exudate, inflammation and microbial contamination; were remarkably decreased upon treatment with collagen gel. On Day 8, a thin tissue layer had formed over the whole surface of the wounds, mostly those treated with collagen gel. At this point, all the lesions had shrunk. Lesions’ size had shrunk by Day 14, and full re-epithelization had been seen in the collagen-based group. When compared to the gel (G3) and the control saline groups, the collagen nanoparticles promoted quicker wound healing and re-hairing. Collagen gel nanoparticles demonstrated improved wound closure due to their ease of application and high adherence to the wound bed.

#### Histopathological analysis

During the healing process, inflammation is the body’s self-defense mechanism for removing essential stimuli such as irritants, pathogens, or damaged cells. Many cells, including macrophages, eosinophils, and neutrophils, participate in the pathogenesis of chronic inflammation by the generation of inflammatory cytokines.

Figs. 9, and 10 show the image of the histopathological examination of the wounded areas of skin from different animals of the treatment groups on Day 10 and Day 21, respectively of H &E stained (X10 & X20). Wounded skins demonstrated a varying response to the treatment, at which the finest wound healing efficacy and full epithelization were verified for the collagen nanoparticles gel G III group followed by the PRP gel GII group. On the 10^th^ and 21^st^ days, the wound region in the treated groups exhibited no ulcer with virtually normal skin covered in dense connective tissues, and a fresh dermal layer.

**Figure (9):**
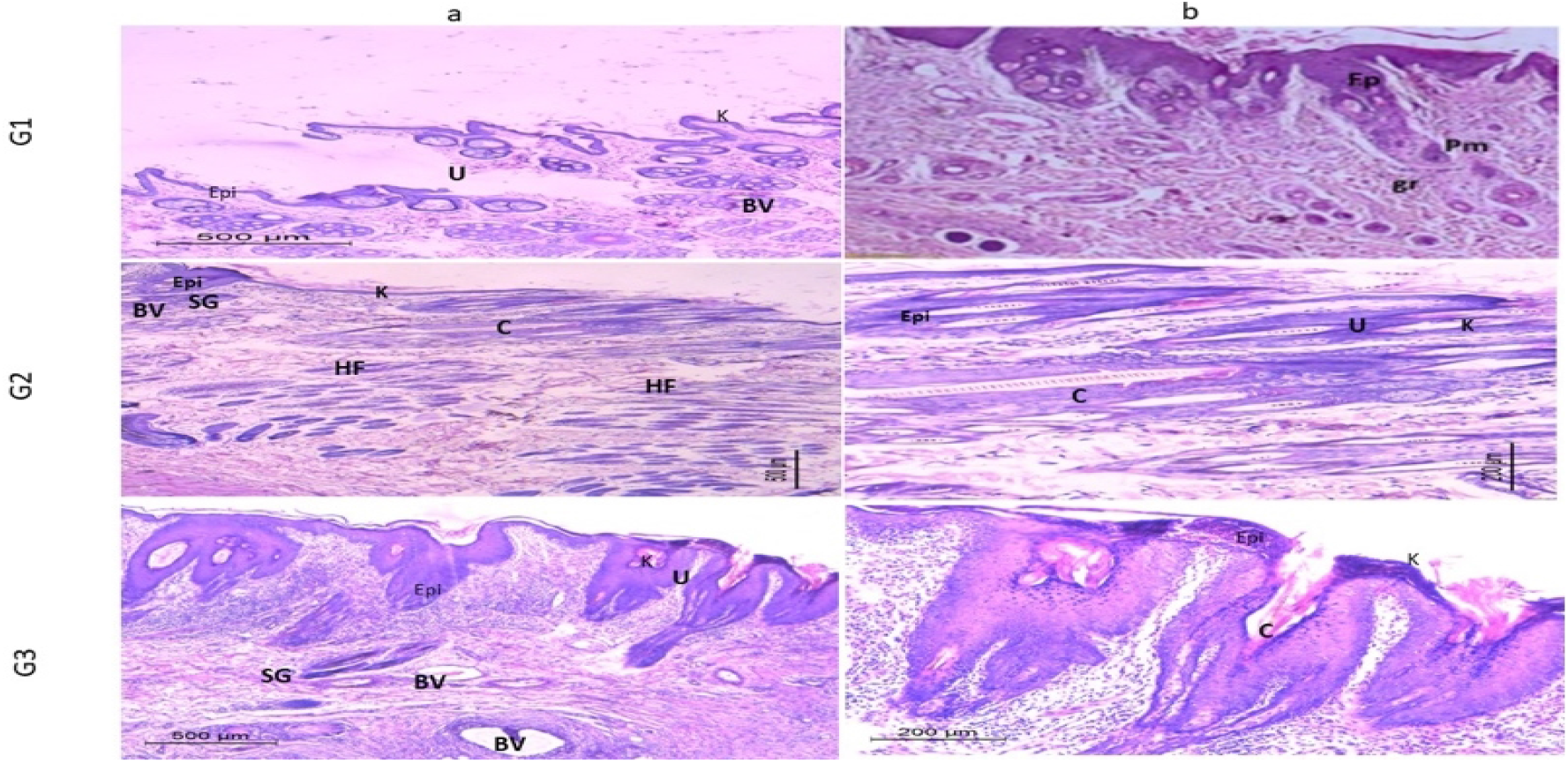
Histological analysis of wounds treated with PRP gel (GII), gel supplemented with collagen nanoparticles (GIII), and saline as negative control (Saline) (GI) at 10th day. Epi: Epidermis; K: keratin layer; BV: blood vessels; HF: hair follicles; SG: sebaceous glands; U: ulcer; C: collagen; Connective tissue (CT) a and b are the images with 10x and 20x magnification.

**Figure (10):**
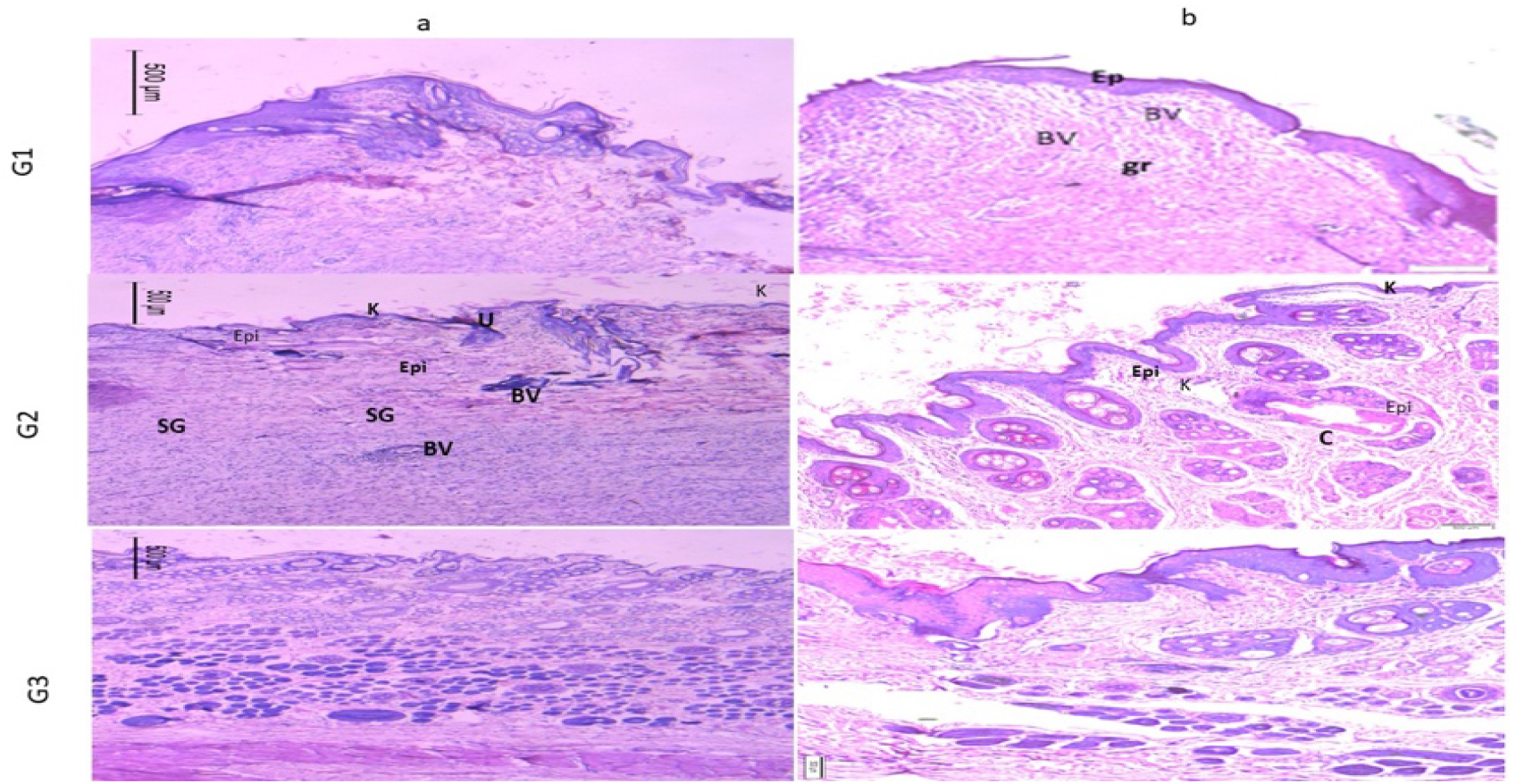
Histological analysis of wounds treated with PRP gel (GII), gel supplemented with collagen nanoparticles (GIII), and saline as negative control (Saline) (GI) at 21st day. Epi: Epidermis; K: keratin layer; BV: blood vessels; HF: hair follicles; SG: sebaceous glands; U: ulcer; C: collagen; Connective tissue (CT) a and b are the images with 10x and 20x magnification.

The image of the histological analysis of the wounds from different treatments groups on Day 10 and Day 21 is shown in Figs. 9 and 10. Animals from the untreated wounds Group GI (Saline group) had ulceration with significant infiltration of inflammatory cells or granulation tissue, but no newly developed blood vessels or hair follicles in H&E stained sections. The dermis was disorganized, with the generation of an inflammatory cap and the presence of a gap, which indicates the incomplete healing of the induced ulcer.

As with typical healed skin, the tissues have a well-formed epidermis (external epithelium made up of 2-3 cell layers) and dermis (connective tissue layer). The most fundamental elements of the rabbit’s skin, such as hair follicles (HF), sebaceous glands (SG), and blood vessels (BV), were also visible in the skin sections. Angiogenesis has been shown to have a critical role in the regeneration of both the epidermis (EPI) and dermis (D) containing hair follicles. The promising action of collagen nanoparticles gel could be attributed to quicker healing and epithelization.

## Discussion

In the current study, we created a medicinal nano-biomaterial from waste collagen and assessed its effectiveness in healing rabbit wounds.Here, we applied the desovlation approach, and as demonstrated by SEM and DLS analysis, the identifiable collagen fibers transformed to accumulations that were spherical in shape (182 nm). These findings are closely related to the research by Jahanban-Esfahlan et al.^18^, who used the desolvation process to produce albumin nanoparticles with a controlled particle size of roughly 100 nm for drug delivery applications. Combining collagen with nanoparticle features (such as better surface-area-to-volume ratio, high porosity, improved mechanical capabilities, and good ability to distribute bioactive compounds) may speed up wound healing and increase skin regeneration^19^.

The bulk of the nanoparticles also had a protein shell on their surface, creating a protein corona that is physiologically active, according to TEM analyses. Depending on the situation, collagen nanoparticle biomedical applications may profit from or suffer from protein corona’s biological impacts^20^. The thin layer of protein corona seems to lessen nanoparticle adhesion and aggregation, inhibiting the identification of macrophages or, in the worst situation, the formation of a thrombus^21, 22^.

The strong negative potential value of collagen nanoparticles (−17.7 mV) as indicated by DLS enhances their long-term stability, superior colloidal nature, and high dispersion due to negative– negative repulsion. ^23^. According to some reports, nanoparticles’ negative potential induces electrostatic repulsions between individual particles, which prevent aggregation. This level of stability is necessary to prevent thrombosis and the aggregation of nanoparticles ^24^.

The created collagen nanoparticles are typically smaller than 200 nm and larger than 5 nm, which makes them a potential drug delivery method for entering cells via endocytosis^25^. Due to electrostatic interactions, it might conveniently attach to the cell membrane as a cationic nanoparticle^26^.

In our research, it was discovered that collagen nanoparticle-infused gel had beneficial impacts on the healing of incision wounds. Our results agreed with those of Mondal et al.^27^, who developed gold-loaded hydroxyapatite collagen nano-biomaterials with improved properties that encouraged cellular adhesion, growth, and proliferation while having bioactive and biocompatibility properties.

*In vivo* testing demonstrated that collagen nanoparticles improved wound healing and produced substantial changes in tissue repair therapies on the tenth day and twenty-first day after damage. Increased cellular proliferation and collagen production at the wound site demonstrated the extraordinary wound-healing ability of collagen nanoparticles. The collagen improvement of wound contraction may have increased the number of myofibroblasts or enhanced their contractile abilities, even though myofibroblasts are regarded to be crucial for the centripetal movement of the wound margin^28^.

The basic objective of a wound dressing, to keep the wound clean and free of external pollutants, appears to be achieved by these nanoparticles^29^. It also keeps the site hydrated, promoting healing and preventing the wound’s origin from being exposed. In addition to being able to transfer active compounds to aid in the healing process, the new nanoparticles have the capacity to protect the wound environment. Compared to the control group, collagen nanoparticles increased wound healing, indicating that they may be employed in wound dressing for skin regeneration.

## Conclusion

In the present investigation, a simple and fast method of desolvation technique was developed for the preparation of nanoparticles from fish scale collagen. The present research highlights novel approaches for wound healing, where collagen nanoparticles derived from fish scales were demonstrated to have qualities similar to type I collagen. Collagen nanoparticles from fish scales are a natural and recycled source of nanoparticles that will be useful in wound healing and other medical applications. Although the expenses are modest since the product is made from natural materials, the most significant consequence is that it will aid in the relief of pain and consequent healing. The main reason for collagen nanoparticles’ promising use in biomedical applications is their nanoscale (172 nm) along with suitable physicochemical properties such as the encircling corona. According to macroscopical and histological analysis, these nanoparticles had connected effectively with the wound region and quickly penetrated the epidermal layer at the wound site because of their nano size and high surface area-to-volume ratio. The synthesized nanoparticles were studied for cell cytotoxicity in the presence of human skin fibroblast cell lines, and no cytotoxic effects were identified for collagen nanoparticles up to 64 µg/ml loading. A little toxicity was observed when the collagen nanoparticles loading reached 1mg/ml. The *in vivo* study of formulation exhibited good wound-healing activity in wound-healing model. Histological studies of healed tissue specify that gel with collagen promotes the wound healing of animal tissue with regenerating hair follicles. The majority of commercially available collagen-based wound dressing treatments are costly and require a secondary dressing. The synthesized and well-characterized collagen nanoparticle system could be cheap and useful for drug delivery as well as suitable extra cellular matrix for tissue engineering and biomedical applications. Unlike our product that was used for 21 days, wound dressings that need to be changed often result in burden and high cost of wound care. Therefore, cost-effective and perfect wound dressings that may be made using cheap and natural products are desperately needed.

## Acknowledgements

This research was funded by the Academy of Scientific Research and Technology (ASRT), Grant number 26D/2017.

## Disclosure

“Manal Shalaby, Ahmed A. Zaki A. Ghareeb, Shaimaa M. Khedr, Haitham M. Mostafa Hesham Saeed and Dalia Hamouda declare that they have no conflict of interest.”

## Compliance with Ethics Guidelines

The study was approved by the Local Ethical Committee of Animal Research, City of Scientific Research and Technological Applications.

## Data availability

All data generated or analyzed during this study are included in this published article.

## Authorship

All named authors meet the International Committee of Medical Journal Editors (ICMJE) criteria for authorship for this article, take responsibility for the integrity of the work as a whole, and have given their approval for this version to be published.”

## Author contributions

All authors contributed to the study’s conception and design. Material preparation, data collection and analysis were performed by [Manal Shalaby], [A. Zaki A. Ghareeb], [Hesham Saeed], [Haitham M. Mostafa], [Dalia Hamouda] and [Shaimaa M. Khedr]. The first draft of the manuscript was written by [Manal Shalaby] and all authors commented on previous versions of the manuscript. All authors read and approved the final manuscript.”

## Notes

### Competing Interest Statement

The authors have declared no competing interest.

